# Desmoplakin is required for epidermal integrity and morphogenesis in the *Xenopus laevis* embryo

**DOI:** 10.1101/464370

**Authors:** Navaneetha Krishnan Bharathan, Amanda J.G. Dickinson

**Author notes:** Corresponding author is Amanda Dickinson.

## Abstract

Desmoplakin (Dsp) is a unique and critical desmosomal protein, however, it is unclear whether this protein and desmosomes themselves are required for epidermal morphogenesis. Using morpholinos or Crispr/Cas9 mutagenesis we decreased the function of Dsp in frog embryos to better understand its role during epidermal development. Dsp morphant and mutant embryos had developmental defects that mimicked what has been reported in mammals. Such defects included epidermal fragility which correlated with reduction in cortical keratin and junctional e-cadherin in the developing epidermis. Dsp protein sequence and expression are also highly similar with mammals and suggest shared function across vertebrates. Most importantly, we also uncovered a novel function for Dsp in the morphogenesis of the epidermis in *X. laevis*. Specifically, Dsp is required during the process of radial intercalation where basally located cells move into the outer epidermal layer. Once inserted these newly intercalated cells expand their apical surface and then they differentiate into specific epidermal cell types. Decreased levels of Dsp resulted in the failure of the radially intercalating cells to expand their apical surface, thereby reducing the number of differentiated multiciliated and secretory cells. Dsp is also required in the development of other ectodermally derived structures such as the mouth, eye and fin that utilize intercalating-like cell movements. We have developed a novel system, in the frog, to demonstrate for the first time that desmosomes not only protect against mechanical stress but are also critical for epidermal morphogenesis.

**Summary Statement:** Critical desmosomal protein, desmoplakin, is required for proper distribution and levels of cytoskeletal elements and e-cadherin. Thus embryos with decreased desmoplakin have defects in epidermal integrity and morphogenesis.

## INTRODUCTION

Desmosomes together with tight and adherens junctions make up the main complexes that provide adhesion between cells. These cell-cell junctions also integrate signals which can regulate a range of cellular processes such as differentiation and cell migration (reviewed in (Delva et al., 2009; Garrod and Chidgey, 2008; Green and Simpson, 2007; Johnson et al., 2014)). But surprisingly, much less is known about the desmosome relative to its other junctional counterparts. We do know that desmosomes provide mechanical strength to tissue, however, additional functions are now being appreciated (Celentano et al., 2017; Dusek et al., 2007; Green and Gaudry, 2000). Moreover, an understanding of the roles of desmosomes in the developing embryo is in their infancy. Therefore, we have utilized the frog, *Xenopus laevis*, as tractable vertebrate model to better define the function of desmosomes specifically during epidermal development.

The importance of desmosomes to the embryo is clear since birth defects caused by mutations to various desmosomal proteins can be devastating and are often lethal in humans (Celentano et al., 2017; Johnson et al., 2014). Further, mice with null mutations in desmosomal proteins die early in development and zebrafish embryos with deficient desmosomal proteins show major developmental defects that would be incompatible with life (Cheng et al., 2005; Gallicano et al., 2001; Gallicano et al., 1998; Goonesinghe et al., 2012). Thus, it is quite likely that desmosomes are indispensable during early embryonic development (Berika and Garrod, 2014). Conditional loss of function of desmosomal components has also revealed important roles in mammalian epidermal development. For example, the ablation of desmoplakin in the epidermis of the mouse embryo revealed fragility, abnormal cell morphologies and disrupted intermediate filament organization (Vasioukhin et al., 2001).

The desmosome, like adherens junctions, are modular structures that both link adjacent cells to each other and to the intracellular cytoskeleton (see reviews such as (Garrod and Chidgey, 2008; Green and Gaudry, 2000; Green and Simpson, 2007)). The transmembrane desmosomal cadherins, consist of desmogleins and desmocollins, and bridge the extracellular space by homophilic binding to complement cadherins on the adjacent cell. These cadherins are anchored in the cytoplasm by the armadillo proteins, plakoglobin and plakophilin. Such armadillo proteins in turn bind to desmoplakin which then forms the critical link to the intermediate filaments such as keratins and microtubules. Desmoplakin is composed of a central rod region that facilitates dimerization, a globular head plakin domain that is required for protein-protein interactions and a C-terminal tail that has three plakin repeat domains, the last of which interacts with intermediate filaments (Delva et al., 2009; Garrod and Chidgey, 2008).

Desmosomes are integral in maintaining tissue integrity and instill mechanical resilience to tissues. This is evident in the severe skin and cardiac defects that arise in autoimmune and genetic diseases where desmosomal components are perturbed (Johnson et al., 2014). Desmosomes can also have non-adhesive functions in the epidermis. For example, the desmosomal protein plakoglobin, has integral functions in cell signaling and can interact with Wnt/beta-catenin (Martin et al., 2009; Yang et al., 2012). Further, desmosomes also regulate the mechanical properties of the cells. For example, loss of desmoplakin function results in decreased cell forces and stiffness’s (Broussard et al., 2017) which could affect cell shape, cell movements as well as mechanotransduction events. Forces across the desmosomal protein desmoglein2, using a tension sensor further suggest that the desmosome is more than a passive rivet like structure (Baddam et al., 2018).

While many of the intriguing functions of desmosomes are unfolding in cells cultured in a dish or in other artificial in vitro systems, it is unclear whether all of these functions also widely apply in vivo, especially during embryonic development. Further how desmosomes might function in cell morphogenesis in an embryo has not been determined. In the present study we focus on a role for desmosomes specifically during epidermal development. In *X. laevis*, the developing epidermis is comprised of an outer or superficial layer and inner or sensorial layer (Billett and Gould, 1971; Drysdale and Elinson, 1992). The outer epidermal layer contains specific differentiated cells including goblet cells, small secretory cells, ionocytes and multiciliated cells. Together these cells protect the embryo from toxins and bacteria by secreting substances and moving fluids (Dubaissi et al., 2014). The inner epidermal layer contains cells with stem cell properties, that is, they continually divide and supply new cells to the outer layer. These inner cells are specified and then move into the outer epidermal layer through a process called radial intercalation (Drysdale and Elinson, 1992; Dubaissi and Papalopulu, 2011; Stubbs et al., 2006; Walck-Shannon and Hardin, 2014). During this process an inner epidermal cell must first move apically and insert between outer epidermal cells at the vertex of 3-4 cells. Then the apical surface of the inner cell expands as it joins the outer layer. Radial intercalation in the epidermis not only provides specific cell types to the outer epidermis but also allows the skin to rapidly expand as the embryo grows. Several of the players regulating epidermal radial intercalation have been defined in Xenopus especially concerning the MCCs (reviewed in (Walck-Shannon and Hardin, 2014)). These cells are first specified by the Delta-Notch activation of lateral inhibition in the inner cell layer (Stubbs et al., 2006). Several transcription factors such as multicilin, Foxj1 and RFX2 are then responsible for the activation of a multitude of genes that are necessary for MCC specification and differentiation (Chung et al., 2012; Stubbs et al., 2008; Stubbs et al., 2012). PCP signals together with microtubule and associated proteins are important for polarizing and inserting the inner cells into the outer epidermal layer (Kim et al., 2012; Kim et al., 2018; Mitchell et al., 2009; Ossipova et al., 2015; Werner et al., 2014). Finally, apical expansion of the intercalating MCC cell is regulated by RhoA signals combined with formins and actin (Sedzinski et al., 2016, 2017). Whether desmosomes have a role in radial intercalation of the epidermis has never been explored.

Development of the mammalian epidermis shares similarities with the developing epidermis of *X. laevis* (reviewed in (Liu et al., 2013)). For example, like in *X. laevis*, the mammalian epidermis begins to develop as two layers. Further, cells from the basal most epidermal layer move apically to form more superficial layers in both mammals and *X. laevis* embryos. The apical movement of basal cells to apical layers is also driven by similar mechanisms in both groups. For example, PCP signals, and microtubule organization are critical in mammalian skin development (Aw et al., 2016; Muroyama and Lechler, 2017). Due to the similarities in epidermal development across vertebrates, we predicted that desmosomes could have a similar roles in the developing epidermis of mammals and *X. laevis*. And if so, Xenopus could serve as a good tool for dissecting the functions of desmosomes in the cell’s biology.

In this study we demonstrate that indeed the desmosome and the desmosomal protein desmoplakin have shared functions, such as providing mechanical integrity, in *X. laevis* and mammals. Importantly, we have developed a novel system, in the frog, to demonstrate that desmosomes are critical for epidermal development.

## METHODS

### *X. laevis* Adults and Embryos

Xenopus laevis adults were created in our breeding colony and purchased from Nasco. All procedures were approved by the VCU Institutional Animal Care and Use Committee (IACUC protocol nμmber 5AD20261). Embryos were collected using standard procedures (Sive et al., 2000) and were staged according to Nieuwkoop and Faber (Nieuwkoop and Faber, 1967). Embryos were cultured in 0.1X MBS, refreshed daily, and housed in a 23°C or 15°C incubator (Torrey Pines Scientific, Cat. No. IN30). After the experiments were completed all embryos were given a lethal dose of anesthetic (10% tricaine for 1 hour).

### Bioinformatics analysis of Dsp

Full-length (Desmoplakin) Dsp protein and mRNA sequences for *Homo sapiens* Desmoplakin I (NP_004406), *Mus musculus* Desmoplakin (NP_076331) and *Danio rerio* Desmoplakin isoform X1 (XP_001919901) *X. laevis laevis* Dsp.L (XB-GENE-866134, Genome Build 9.1, http://www.xenbase.org), were aligned using the LALIGN tool (EMBL-EBI) and the EMBOSS Water tool (Smith Waterman algorithm) (EMBL-EBI) was used to determine similarity. Protein domains were based on those identified in the hμman Dsp protein (Al-Jassar et al., 2011; Green et al., 1990; Virata et al., 1992).

### TEM to examine desmosome ultrastructure and localization in the epidermis

Embryos were processed for EM at the VCU miscoscopy core using standard protocols. Briefly, embryos were fixed with 2% glutaraldehyde (MP Biomedicals, 198595) in 0.1M sodiμm cacodylate buffer (Electron Microscopy Services, 12300) overnight (4°C) and then refixed in 2% osmiμm tetroxide in 0.1M cacodylate buffer (one hour). They were dehydrated in ethanol and then infiltrated withpropylene oxide (EMS, 20401) and Poly/Bed 812 resin mix (50:50, Polyscienes, 08792-1) overnight. Finally embryos were incubated in EMbed 812 resin (EMS, 14120, overnight), placed in molds and heated 60°C (2 days). 700-900Å thick sections were created on a Leica EM UC6i Ultramicrotome (Leica Microsystems) and stained with 5% Uranyl acetate (EMS, 22400) and Reynold’s Lead Citrate (Lead Nitrate (EMS, 17900) and Sodiμm Citrate (EMS, 21140)). A JEOL JEM-1230 TEM (JEOL USA, Inc.) with the Gatan Orius SC1000 digital camera (Gatan Inc., Pleasanton, CA) was used to image the ultrastructure of the epidermis.

### Dsp Morpholinos and RT-PCR tests for splicing defects

Functional analysis was performed utilizing splice blocking antisense Morpholinos (Genetools). DspMO1 (5′-ACAGTTACTACTTACTCTATGCTGC-3′) targets donor site of exon 4 and DspMO2(5′–TTGATGCAGAGCAAAGTTCAAACCT-3′) targets the donor site of exon 14. The standard control morpholino (CMO) was used in all experiments at 34-68ng/embryo. Morpholinos were labeled with fluorescein to gauge injection success and for blastomere targeting. A FemtoJet microinjector (Eppendorf) and a SteREO Discovery.V8 (Zeiss) stereoscope were used for microinjections. Embryos injected at the 1 cell stage were injected with 17-35 ng while those injected at the 16 cell stage were injected with 1-3 ng of Dsp morpholinos.

RT-PCR was performed as described previously (Dickinson and Sive, 2009) to test for splicing defects. RNA was extracted with TRIzol (invitrogen) followed by lithiμm chloride precipitation. cDNA was prepared using the High-capacity cDNA Reverse Transcription kit (Applied Biosystems). Dsp primers flanking the targeted exons (provided upon request) were used together with the Apex™ Hot Start Taq DNA Polymerase Master Mix.

### Dsp CRISPR/Cas9 mutational analysis

Injectable gRNA was created as described previously (Shah et al., 2016). CHOPCHOP (http://chopchop.cbu.uib.no/) was used to design two CRISPR gRNA sequences with no mismatches and high efficiency targetingexons 8 (dspCrispr1) and 19 (dspCrispr2) of*dsp.L* (these correspond to exons 8 and 17 in *dsp.S*, respectively). The gRNA sequences were GGTGCTGGTTCATGATAAGC<u>**TGG**</u>(DspCrispr1) GGTCGCATCTGACAGTTTGA<u>**TGG**</u> (DspCrispr2).A T7 primer sequence and loop-specific sequence (5′-AATTAATACGACTCACTATA-(N)_20_-GTTTTAGAGCTAGAAATAGC-3’) were added to create a DspCrispr Oligo (ordered from IDT). Double stranded DNA template was produced combining the customs designed oligo with a generic Scaffold Oligo (5′GATCCGCACCGACTCGGTGCCACTTTTTCAAGTTGATAACGGACTAGCCTTATTTTAACTTGCTATTTCTAGCTCTAAAAC-3′; IDT) and PCR performed with the following parameters; 98°C (30s), 98°C (10s), 61°C (10s), 72°C (15s) X 45 cycles, followed by a 72°C (5 min) extension. DNA was then processed through the Clean & Concentrator Kit (Zymo Research, D4014) to purify and concentrate the template. The Ambion MEGAscript T7 kit (Thermo Fisher Scientific, AM1333) was used for *in vitro* transcription following the manufacter’s instructions with long incubation (16hrs-24hrs). To create F0 mosaic mutants, embryos were co-injected with 1ng gRNA and 1.5ng Cas9 protein (1mg/ml PNA-Bio, CP01) at the one-cell stage. Wild-type embryos were used as controls.

The T7 endonuclease I assay was used to detect mutations induced by CRISPR/ Cas9 as described previously (Mashal et al., 1995). Embryos were incubated in a lysis buffer consisting of 25 mM NaOH (Fisher Scientific, BP359) and 0.2 mM Na2+-EDTA (OmniPur, 4050)) at 95°C for 40 minutes. After the samples cooled an equal volμme of neutralization buffer (40 mM Tris-HCl (Sigma-Aldrich, T3253)) was added. PCR was performed with 2X Phusion master mix and primers flanking the predicted mutation site (Primers available upon request), using the following parameters 98°C (30s), 98°C (5s), 61°C (10s), 72°C (20s) for 36 cycles with a 72°C (2min) extension. PCR products were purified with the DNA Clean and Concentrator kit (Zymo Research, D4014), eluted into nuclease-free water, and quantified using the NanoDrop Lite spectrophotometer (Thermo Fisher Scientific). Purified PCR product (200ng) was used in the following protocol: 95°C (5 min.), 95°C-85°C (−2°C/s), 85°C-25°C(−0.1°C/s). Then, 1μl of T7 endonuclease I (NEB, M0302) was added and the solution incubated at 37°C for 15 min. Samples were run on a 3% agarose gel to identify the presence of mutations.

### Mechanical stress assays

#### Dropping Assay

Embryos were vertically dropped from a height of 15 cm using a transfer pipette (Fisher Scientific, 13-711-7M) onto a 150 × 15mm petri dish lined on the bottom with 5mm of 2% agarose (Bioline, 41025). 0.1XMBS was added to the dish to submerge the embryo and then it was visualized and photographed. Images were taken on both left and right lateral sides of the embryos before and after the assay was performed. This procedure was done blindly to avoid any handling bias while pipetting embryos.

#### Rotational Assay

Embryos were placed in 50ml plastic polypropylene tubes (USA Scientific, 1500-1811) with 15 ml 0.1 X MBS (Modified Barth’s Saline). The tubes were then rotated using a RKVSD vertical rotating mixer (ATR Biotech) at 55 rpm for a total of 25 rotations.

##### Quantification and Statistical Analysis

Embryos were scored as either damaged or undamaged based on whether the epidermis was visually intact or not. Chi-Squared tests were performed in Excel to determine statistical relationships between treatment and control groups. Error bars representing standard error were also calculated in Excel.

### Histology

Histology was performed as described (Dickinson and Sive, 2006; Houssin et al., 2017) with some modifications. Briefly, embryos were fixed in 2% PFA and 2% glutaraldehyde in PBT buffer for 24 hr and then embedded in plastic resin (JB-4 Plus) and sectioned at 5-7 μm using a tungsten carbide knife. Sections were stained with Giemsa at 1:20 for 1 hr followed by 10 sec 0.05% acetic acid differentiation wash. Slides were dried and covered with Permount and imaged on a Nikon compound microscope fitted with a digital camera (VCU Biology microscopy core).

### Immunohistochemistry, Phalloidin, PNA

Embryos were fixed in 4% Paraformaldehyde (PFA) or Dent’s fixative (80% methanol:20% DMSO) and then labeled whole or after vibratome sectioning. For sectioning, embryos were embedded in 5% low-melt agarose (SeaPlaque GTG Cambrex) and sectioned using a 5000 Series Vibratome into 150-200μm sections. Primary antibodies included; mouse anti-desmoplakin I+II (abcam, ab16434, diluted 1:75), mouse anti α-tubulin (Developmental Studies Hybridoma Bank (DSHB), AA4.3, 1:50), mouse anti-cytokeratin type II (DSHB, 1h5, 1:25), anti-β-catenin (Invitrogen, 71-2700, 1:500), and mouse anti-E-cadherin extracellular domain (DSHB, 5D3, 1:25). Dsp was detected using tyramide amplification (Alexa Fluor 488 Tyramide Superboost, goat anti-mouse, Invitrogen b40941). All other antibodies were detected with anti-mouse or anti-rabbit Alexa Fluor’s used at 1:500 (Invitrogen). Fluorescently labeled substrates included Lectin PNA (Alexa Fluor 488, Invitrogen, L21409, 1:1000) and phalloidin (Rhodamine,Life Technologies, R415, 1:50).

### Relative keratin intensity profiles

Keratin intensity profiles were generated from images acquired from a C2 Nikon confocal microscope. In the accompanying Nikon Elements software Intensity profile lines (white lines with an arrow) were added to the image so that they crossed beta-catenin labeling at cell junctions and extended into the cytoplasm of each cell. The pixel intensity of beta-catenin and keratin labeling along the intensity profile line was plotted in Elements software.

### Biotin labelling

Epidermal surface labeling was achieved by incubating embryos for 1 hour in EZ-Link Sulfo-NHS-LC-Biotin (10mg/ml; Thermo Fisher Scientific, 21335) dissolved in 0.1X MBS. This was followed by a 10μm glycine wash for 10 minutes and then a washout period of 4 hours in 0.1XMBS. Embryos were fixed in 4% PFA overnight at 4°C, blocked with 1% BSA and then labelled with Streptavidin conjugated to Alexa Fluor 568 (Life Technologies, S11226, 1:500). Biotin labeling was imaged with a C2 Nikon confocal.

### Quantification of surface areas and nμmbers of cells

In all quantification experiments, images were taken of the lateral trunk midway between the head and tail tip using the 20X objective on a C2 Nikon confocal.

The relative surface areas of MCC cells labeled with anti-tubulin and phalloidin were measured using the magic wand tool in Photoshop. The wand was drawn along the outside of the cortical actin labeling to estimate cell surface area. Ten cells were measured in each confocal image field. A total of 10 embryos were sampled from three different experiments. The surface areas of the biotin negative cells were similarly measured using the magic wand tool in Photoshop by outlining the border of the cell at the junction of labeling and unlabeling. All cells in each confocal image field were measured. <u>Statistical Analyses:</u> Sigmastat software was utilized to determine differences between Dsp morphants and control groups. Normality tests were first performed using Shapiro-Wilks test. In the Biotin surface analysis the normality test failed and therefore a Mann-Whitney Rank Sum Test was performed. In the analysis of surface areas of tubulin positive cells the normality test passed and a student t-test was performed. Sigmastat or Sigmaplot was used to create box and whisker plots where the 5^th^ and 95^th^ interval are shown and median and mean lines provided. Histograms were created with equal binning and automated scaling adjusted for equivalent comparisons.

The number of tubulin positive cells, PNA positive regions or cells unlabeled with biotin were quantified by counting all cells (including unlabeled cells) in each image field to achieve relative quantities. Image fields taken from 10-11 embryos (from at least 2 biological replicates) were quantified. <u>Statistical Analyses:</u> SigmaStat software or Excel was utilized to determine differences between Dsp morphants and control groups as well as to create box and whisker plots or bar graphs. Normality tests passed in all analyses and therefore student t tests were utilized.

### Confocal imaging and Photoshop processing

Embryos were imaged in 90-100% glycerol in PBT using a Nikon C2 confocal microscope located in the Biology Department Microscopy Core. Z-stacks were created with a step size of 0.3-0.5 μm, for a total thickness between 2–15 μm. Using NIS-Elements AR 4.50.00 software, Z-stacks were converted to maximum projection images depending on the experiment. Adobe Photoshop was used to further process images, which include increasing the brightness, cropping and labeling. Changes to the channel color were performed to ensure consistency and accommodate those with color blindness. All images for each experiment were processed in the same way.

### DP-NTP expression in embryos

DP-NTP, created by Dr. Kathleen Green, was cloned into the pCS2+ plasmid by Dr. Daniel Conway (Department of Biomedical Engineering, VCU) and provided as a kind gift. To create mRNA, the plasmid was digested with HpaI and then was transcribed using SP6 mMessage mMachine kit (Ambion). 1 ng was injected into embryos at the one cell stage.

## RESULTS

### 1. Desmosomes and desmoplakin exist in the developing epidermis

Our first goal was to characterize desmosomes in the developing epidermis of *X. laevis*. In this species, this epidermal tissue is composed of two layers; an inner and an outer layer (Fig. 1A). We utilized a combination of transmission electron microscopy (TEM) and immunofluorescence to examine desmosomes and desmoplakin (Dsp) the inner and outer epidermal layers during development.

**Figure 1:**
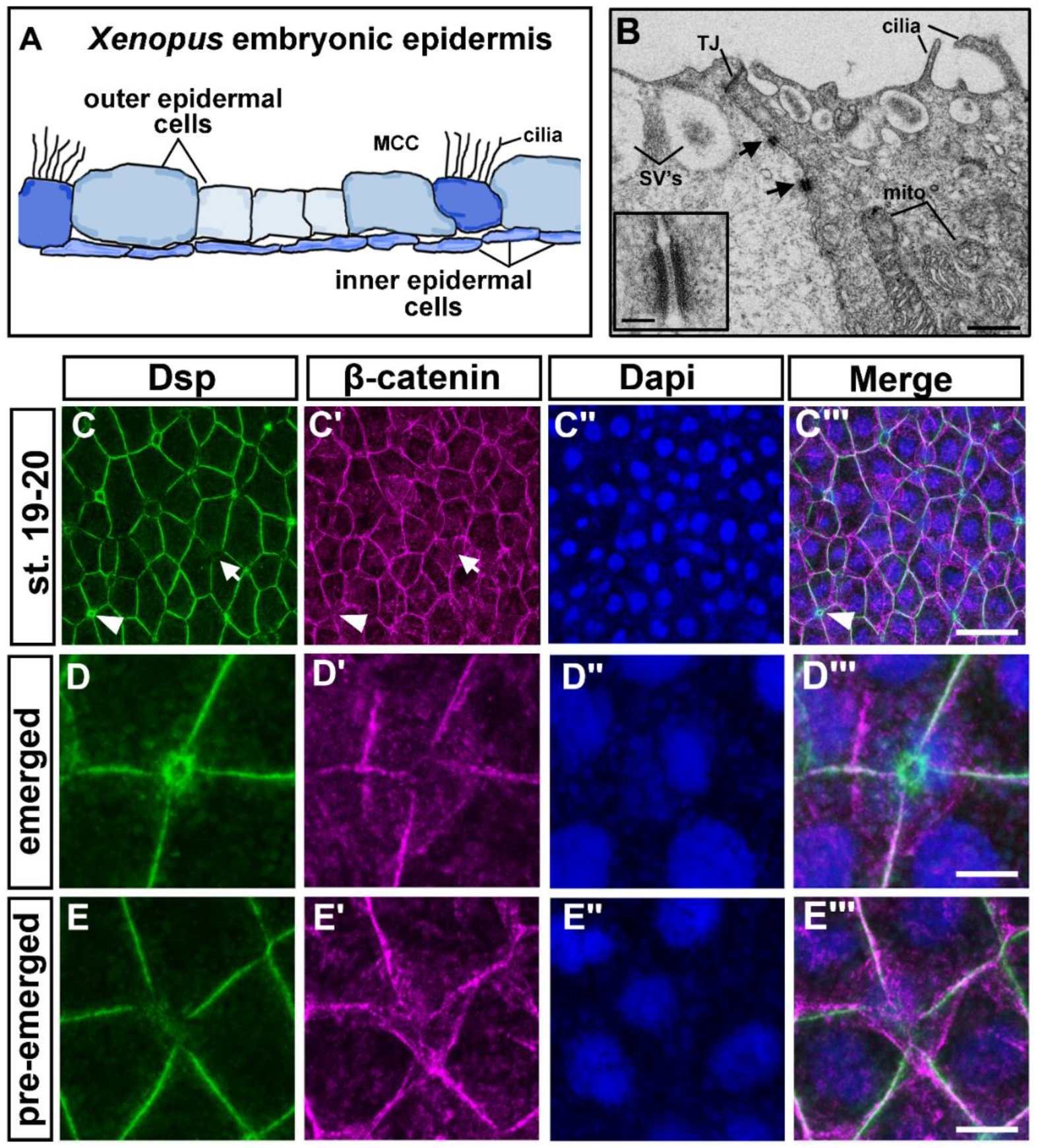
Desmosomes and desmoplakin in the epidermis of *X. laevis* embryos. A) Schematic showing a transverse section of the *X. laevis* embryonic epidermis based on actual histological sections. B) TEM images where black arrows indicate desmosomes located between outer epidermal cells. Scale bar=1.8μm. Inset shows higher magnification of a desmosome. Scale bar= 135nm. C-D‴) Immunofluorescence of Dsp and beta-catenin in the epidermis. Dapi labeling of the nuclei was used as a counter stain. C-C‴) st. 19-20, in H arrows indicate regions where there is a less Dsp labeling and arrowheads indicate where there is enrichment of Dsp labeling. The same cells are indicated in panel C′. Scale bar=27μm. D-D‴) High magnification of a cell that appears to be apically emerging with high levels of Dsp and lower beta-catenin. Scale bar = 6μm. D-D‴) High magnification of a cell that appears to be beginning to intercalate and has low levels of Dsp. Scale bar=6μm. Abbreviations= MCC= multiciliated cell, SVs=secretory vesicles, mito=mitochondria, TJ=tight junction, oe=outer epidermis, ie=inner epidermis.

#### 1a) TEM reveals that desmosomes are present in the epidermis during embryonic development of X. laevis

The location of desmosomes in the epidermal layers was determined using TEM. Desmosomes were observed frequently at junctions between outer epidermal cells at all stages examined ranging from stage 10 (9hpf) to stage 45 (96hpf)(Fig. 1B and Fig. S1). Desmosomes were always observed below a tight junction and could be observed between different cell types (Fig. 1B, shows ciliated cell adjacent to a secretory cell). At higher magnification, the ultrastructure of *X. laevis* embryonic desmosomes resembled desmosomes described in other vertebrates and a subset contained a classic dense intercellular midline (Borysenko and Revel, 1973; Garrod et al., 2005) (Fig. 1B inset). We also observed desmosomes to have connections to a filamentous network consistent with intermediate filaments (Fig. S1A). These complexes were also infrequently detected between outer and inner epidermal cells (Fig. S1B-C, black arrowheads) as well as between the inner epidermal cells (Fig. S1B-C, white arrowhead). Thus, desmosome ultrastructure, position in the cell and the connections they make in the developing frog embryo are remarkably similar to what has been reported in mammals.

#### 1b) Desmoplakin is a conserved protein that is expressed in the epidermis of the developing X. laevis embryo

Desmoplakin (Dsp) is unique to the desmosome and the primary connection to both the cytoskeleton and the desmosomal cadherins. Therefore, this protein serves as a good proxy to understand the development and function of desmosomes in the *X. laevis* embryo. To support this idea, *dsp* is expressed throughout development as assessed by RT-PCR (Fig. S2A) which is consistent with published RNAseq data (Session et al., 2016). Sequence alignments revealed that the *X. laevis* Dsp mRNA and protein are also highly similar to human and mouse (Table 1). For example, the *X. laevis* Dsp protein is 82% similar to the hμman Dsp protein. Additionally, the frog Dsp protein has all the same functional domains as the human Dsp protein unlike zebrafish (Table 1). Finally, the high level of sequence similarity in the Dsp protein was not observed in bioinformatic comparisons of other desmosomal proteins (not shown). Taken together, this data supports the idea that *X. laevis* Dsp is an ideal protein to examine and perturb to study the role of desmosomes during epidermal development.

**Table 1.**
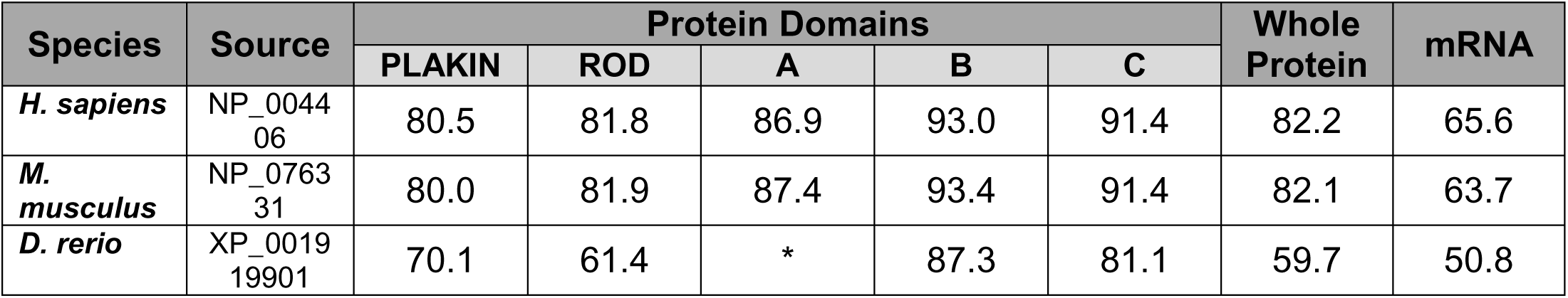
Comparative analysis of *X. laevis Dsp.L* desmoplakin mRNA and protein sequences with human, mouse and zebrafish. % Similarity is provided. Sequences not present or available is indicated by an asterisk. Xenbase was used to access genome builds for *X. laevis*.

| Species | Source | Protein Domains |  |  |  |  | Whole Protein | mRNA |
| --- | --- | --- | --- | --- | --- | --- | --- | --- |
|  |  | PLAKIN | ROD | A | B | C |  |  |
| *H. sapiens* | NP_004406 | 80.5 | 81.8 | 86.9 | 93.0 | 91.4 | 82.2 | 65.6 |
| *M. musculus* | NP_076331 | 80.0 | 81.9 | 87.4 | 93.4 | 91.4 | 82.1 | 63.7 |
| *D. rerio* | XP_001919901 | 70.1 | 61.4 | * | 87.3 | 81.1 | 59.7 | 50.8 |

Immunofluorescence revealed Dsp protein in the outer ectoderm and epidermis, from gastrulation to early tadpole stages (Fig. 1C-E‴, Fig. S2B-E‴). Beta-catenin and Dapi were used as a counterstain in our imaging of Dsp to better visualize the developing epidermis. During gastrulation, Dsp protein was localized in a discontinuous pattern adjacent to beta-catenin in the outer epidermal cells (Fig. S2 B-B‴). At stages 19-20 (20-22 hpf), when morphogenesis and differentiation of the epidermis begins, we noted that Dsp was not equal in intensity in all outer epidermal cells (Fig. 1C-C‴). For example, some cells with small apical surface areas appeared to be enriched with Dsp while beta-catenin was depleted in the same region of these cells (Fig. 1C-C‴, arrowheads, Fig. 1D-D‴, suppl. movie 1). Such cells with small apical surfaces are consistent with newly emerged intercalated cells. At the vertices of 4-5 cells, where cells were in the process of inserting in the outer ectoderm, Dsp levels appeared lower (Fig. 1C-C‴, arrows, Fig. 1E-E‴ and suppl. movie 2). Later, at stages 24-25 (24-27 hpf) and 32-33 (40-45 hpf), Dsp appeared to be more evenly distributed in the outer epidermal cells (Fig. S2D-E‴) and could be visualized in a punctuate pattern in the inner epidermal cells at lower intensities (Fig. S2F-G‴, arrows).

These results demonstrate that Dsp was present in all cells of the developing epidermis and differences in localization are evident in cells undergoing radial intercalation.

### 2. Morpholinos and Crispr/Cas9 mediated depletion of Dsp reveal its importance in *X. laevis* embryonic development

We demonstrated that desmosomes and the major desmosomal component, desmoplakin (Dsp), are present in the epidermis of the embryo. To better determine the specific functions of desmosomes and Dsp during epidermal development and morphogenesis morpholino (MO) and Crispr/cas9 technologies were utilized to deplete Dsp.

#### 2a) Morpholinos and Crispr/Cas9 technologies results in Dsp reduction

Since MOs are titrate-able, these can be utilized to examine the effects of decreased dosage of Dsp in the embryo. This is especially useful when a complete knockout might be lethal as Dsp is in the mouse (Gallicano et al., 1998). Two antisense splice-blocking MOs targeted to the splice donor sites of exon 4 (DspMO1) and exon 14 (DspMO2) (Fig. 2A, B) were utilized. Various amounts of MO were tested until a consistent and effective amounts were established. To determine the efficacy of the morpholinos, morphants injected with 34ng/embryo DspMO1 and 17ng/embryo (DspMO2) were analyzed for splicing defects using RT-PCR as well as Dsp protein reduction using immunofluorescence. Results revealed that indeed DspMO1 and DspMO2 injections resulted in splicing defects (n=10, 2 experiments, Fig. 2A, B) and a reduction in desmoplakin protein (n=10, 2 experiments, Fig. 2A,B).

**Figure 2:**
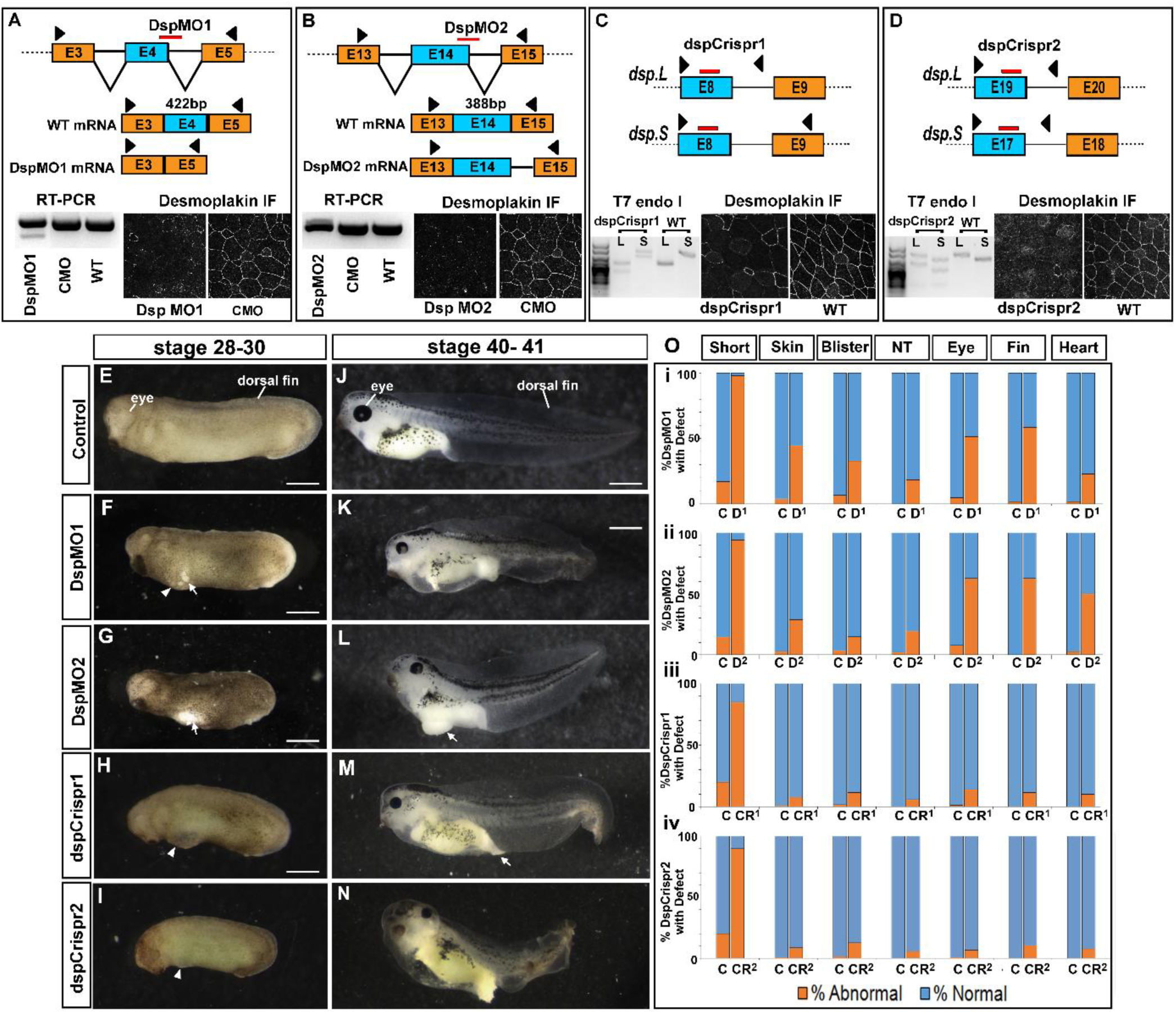
Decreased function of Dsp in the embryo.(A) Schematic showing where the DspMO1 (red bar) is predicted to bind to *dsp.L* mRNA as well as the predicted spliced products in Dsp morphants and controls. Black arrow heads indicate the region of primer sequences used for RT-PCR analysis. RT-PCR of *dsp* showing there is an alternative splice product in the DspMO1 morphants. Immunofluorescence of desmoplakin (white) in DspMO1 and CMO morphants. (B) Schematic showing where the DspMO2 (red bar) is predicted to bind to *dsp.L* mRNA as well as the predictedspliced products in Dsp morphants and controls. Black arrow heads indicate the region of primer sequences used for RT-PCR analysis. RT-PCR of *dsp* showing there is an alternative splice product in the DspMO1 morphants. Immunofluorescence of desmoplakin (white) in DspMO2 and CMO morphants. (C) Schematic showing where the dspCrispr1 (red bars) is predicted to target the *dsp.L* and *dsp.S* genes. Black arrow heads indicate the region of primer sequences used for mutation analysis. Results of a T7 endonuclease assay are shown of dspCrispr mutants showing there is an alternative product indicative of mutation. Immunofluorescence of desmoplakin (white) in dspCrispr mutants and controls. (D) Schematic showing where the dspCrispr2 (red bars) is predicted to target the*dsp.L* and *dsp.S* genes. Black arrow heads indicate the region of primer sequences used for mutation analysis. Results of a T7 endonuclease assay are shown of dspCrispr mutants showing there is an alternative product indicative of mutation. Immunofluorescence of desmoplakin (white) in dspCrispr mutants and controls. E-I) Lateral view of control, Dsp morphants and mutant embryos at stage 28-30. Anterior is to the left. Arrows indicate regions where the epidermis is broken and arrowheads point to blister like structures. Scale bars=450μm. J-N) Later view of control, Dsp morphants and mutant embryos at stage 40-41. Anterior is to the left. Arrows indicate regions where the epidermis is broken and arrowheads point to blister like structures. Scale bars=450μm. O) Analaysis of phenotypic frequency in i) DspMO1 morphants (D1), ii) DspMO2 morphants (D2), iii) dspCrispr1 mutants (Cr1), dspCrispr2 mutants (Cr2) compared to controls (C). Orange indicates percentage with the defect indicated at the top. Abbreviations: WT=wildtype uninjected sibling embryos, E=exon, IF=immunofluorescence.

While morpholinos are an effective method to test the role of gene function in *X. laevis*, non-specific effects are possible (Blum et al., 2015; Eisen and Smith, 2008; Heasman, 2002). Therefore, to ensure that the phenotypes produced by the Dsp MOs are specific to Dsp reduction, we also utilized the CRISPR/Cas9 system to mutate *dsp* (Bhattacharya et al., 2015; Nakayama et al., 2013; Wang et al., 2015). Two different mosaic F0 *dsp* CRISPR mutants were created where some cells contained the mutation while others did not. To do this, gRNAs targeting exons of the dsp gene (dspCrispr1, Fig. 2C and dspCrispr2, Fig. 2D) were injected with Cas9 protein into embryos. To determine the efficacy of this technique, a subset of injected embryos (displaying a phenotype) were analyzed for mutations using T7 endonuclease assay. Results revealed that indeed prospective dspCrispr1 and dspCrispr2 mutant embryos were positive for regions of non-perfectly matched DNA as predicted (n=12, 2 experiments for each, Fig. 2C,D). To test whether these mutations resulted in a reduction in Dsp protein, immunofluorescence was utilized. dspCrispr1 and dspCrispr2 mutant embryos had regions where cells displayed a clear reduction in Dsp protein (n=12, 2 experiments for each, Fig. 2C).

In summary, these results together demonstrate that morpholino and Crispr/Cas9 methods caused splicing defects and mutations respectively, which in turn effectively reduced Dsp protein in developing embryos.

#### 2b) Decreased Dsp using morpholinos or Crisprs results in identical developmental abnormalities

The defects observed in the DspMO morphants and the dspCrispr F0 mutants were remarkably similar (Fig. 2E-N). For example, at tailbud stages (st. 24-28) the Dsp morphant and mutant embryos were smaller and had epidermal tears, hyperpigmentation, ventral blisters and neural tube closure defects (Fig. 2 E-I, O). Later, at larval stages (st. 40-41), many morphant and some mutant embryos were more likely to have craniofacial defects, smaller eye as well as heart edema and ruffled tail fins (Fig. 2 J-N, O). The percentage of embryos, (at both stages) with these phenotypes was lower in the dspCrispr injected embryos (Fig. 2O). It is possible that the mosaic F0 dspCripsr mutants have such reduced defects because the phenotypes are caused by a large region of epidermal depletion of Dsp in the morphants. The interspersion of normal cells in dspCrispr mutants may prevent the epidermal tearing for example.

In summary, these results suggest that depletion of desmoplakin is required for embryonic growth, epidermal development and for the formation of the face, heart and fin. Importantly, the striking similarities in the Dsp morphants and dspCrispr mutants suggest that these abnormal phenotypes are specifically caused by decreased in Dsp.

### 3. Dsp is specifically required for epidermal integrity and resilience

Since desmosomes are well known for their role in providing mechanical resiliance to the epidermis in mammals we next tested whether this was also true in *X. laevis* embryos. Quantitative assays were developed to test whether the epidermis is more susceptible to mechanical insults when depleted of Dsp. Further, specific loss of Dsp in the epidermis was tested and skin fragility observed.

#### 3a) Dsp morphants are more susceptible to mechanical stress

One explanation for increased epidermal tears in the Dsp morphant and mutant embryos is that they are more fragile and thus less resistant to mechanical perturbations. Here, we developed two methods to quantify the effects of mechanical insults to the epidermis of the Dsp morphants.

In the “Dropping Assay”, embryos were dropped from a height of 15 cm into a dish lined with agarose (Fig. 3A and suppl. movie 3). We expected that this assay could cause shear and impact stresses as the embryo fell and contacted the surface tension of the media. Results demonstrated that indeed significantly more DspMO1 morphants (85%) had damage to the epidermis compared to controls (15%)(Chi-squared test, p=9.55E-06; Fig. 3A-C).

**Figure 3:**
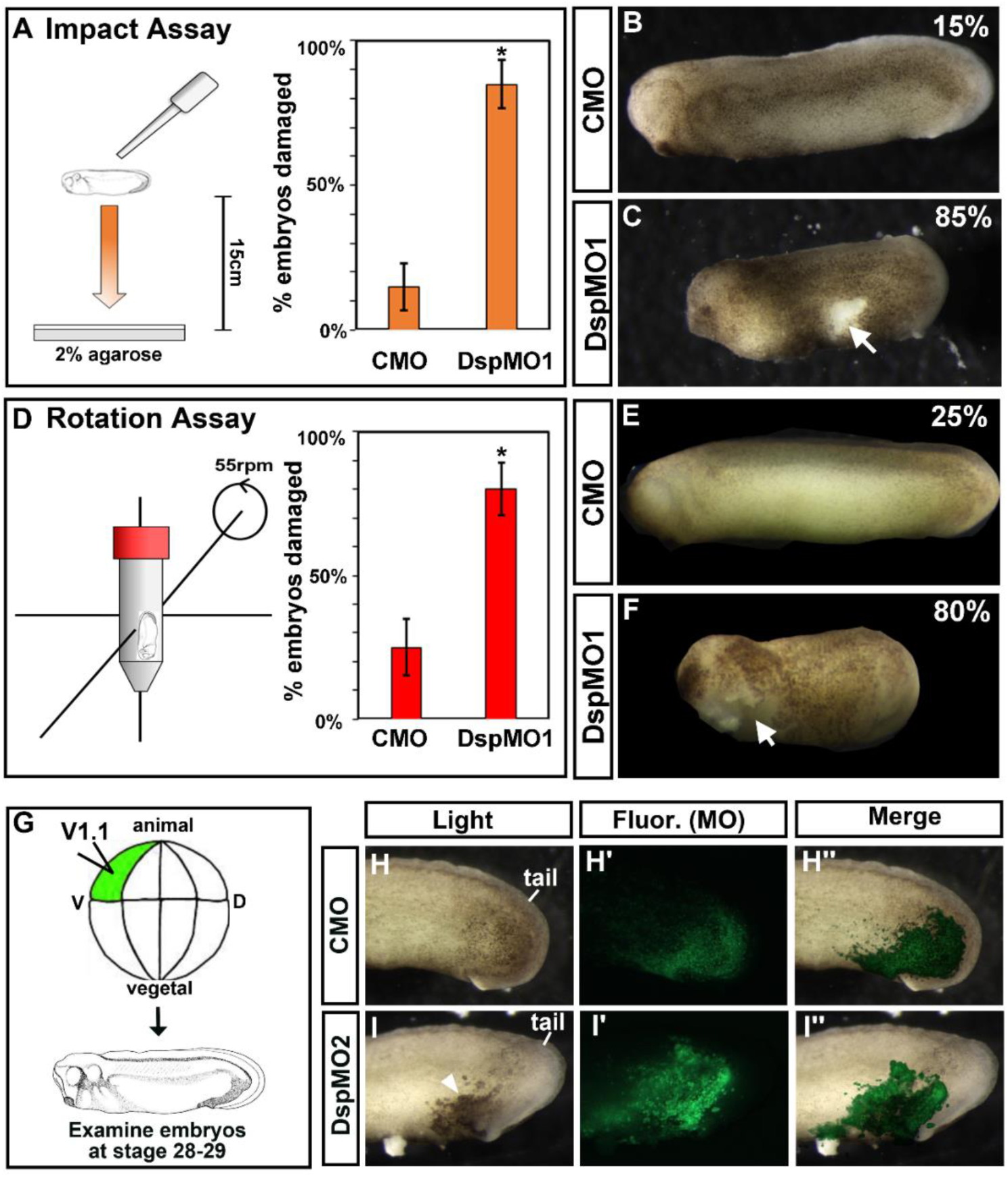
Decreased Dsp results in less mechanical resilience and is specific to the epidermis.A) Schematic of experimental design for the Dropping Assay. Bar graphs summarizing quantification of the proportion of CMO and DspMO1 morphants with a ruptured epidermis after undergoing the Dropping Assay.*=statistical significance. B-C) Lateral views of a representative control and Dsp morphant at stage 28-30. Anterior is to the left. Arrow indicates regions where the epidermis is broken and arrowheads point to blister like structures. Scale bars=450μm. D) Schematic of experimental design for the Rotation Assay. Bar graphs sμmmarizing quantification of the proportion of CMO and DspMO1 morphants with a ruptured epidermis after undergoing the Rotation Assay. *=statistical significance. E-F) Lateral views of a representative control and Dsp morphant at stage 28-30. Anterior is to the left. Arrow indicates regions where the epidermis is broken and arrowheads point to blister like structures. Scale bars=450μm. G) Schematic of epidermal target injections. H-I‴) Representative lateral views of the posterior of embryos as in G showing the embryo under light (Light), fluorescence (Fluor. MO) to visualize the morpholino alone and merged (Merge). Scale bars=450μm. H-H‴) Control MO (CMO) I-I‴) DspMO1 morphants.

In the “Rotational Assay”, embryos were placed in a buffer-containing 50ml conical tube that underwent rotations in a vertical rotating mixer (Fig. 3D and suppl. movie 4). The embryos were likely subject to multiple mechanical stresses as they contacted the surface tension of the media as well as the sides of the tube. Results demonstrated that again significantly more DspMO1 morphants (80%) had damage to the epidermis compared to controls (25%) (Chi-squared test, p=0.000478; Fig. 3 D-F).

These assays demonstrate that the epidermis of DspMO1 morphants have quantitatively decreased resistance to mechanical stress such as impact, shear and/or tensional forces.

#### 3b) Dsp targeted to the trunk epidermis reveals a specific requirement in this tissue

We next tested whether the epidermal integrity defects caused by the Dsp MO were due to a specific requirement for Dsp in the epidermis. It is possible that the epidermal integrity defects observed in Dsp morphants and mutants are due to secondary effects generated from abnormal gastrulation or smaller size of the embryo. To circumvent such non-specific possibilities, targeted injections of fluorescently-tagged DspMO1 were performed. This allowed us to investigate the effects of depleting a subset of epidermal cells of the Dsp protein in the context of a relatively normal embryo. DspMO1 or CMO was injected into a ventrally located blastomere fated to become the trunk epidermis (D1.1; Xenbase, Fig 3G). Results revealed that DspMO1 positive epidermal regions formed tears or ruptures in 100% of the embryos. Such tears were never observed in CMO injected embryos (n=30, 3 experiments, Fig 3B-C″). These results reveal that indeed epidermal integrity is due to Dsp depletion in the epidermis and not due to defects in other tissues.

#### 3c) Dsp depletion results in perturbed desmosomes and increased intercellular space in the outer epidermis, which could account for epidermal integrity problems

Next we aimed to explore the cellular perturbations in DspMO1 morphants that could help account for the increased epidermal fragility. First, we were interested in how the ultrastructure of the outer epidermis was affected by reducing Dsp. In these experiments, embryos were injected at the 1-cell stage and examined at stages ranging from 26-28 (30-35 hpf). TEM demonstrated that desmosomes were often missing in the Dsp morphants (observations made from 50 images taken from 2 embryos, Fig. 4A-D). Missing desmosomes were rarely observed in the control embryos at this stage (2 embryos, 50 images). In addition, at junctions where desmosomes were absent in the Dsp morphants, a large intercellular gap was often observed below an intact tight and/or adherens junction (Fig. 4C,D, arrowheads). Such gaps could be a sheering artifact of the processing due to the lack of integrity in the epidermis. On the other hand they could reflect a change in the extracellular space with the loss of desmosomes. Regardless, together these observations suggest that desmoplakin is required for maintaining desmosomes in the developing epidermis which in turn could be required to maintain strong cell to cell connections. A weakening of cell-cell connections could account for the lack of resilience in the epidermis.

**Figure 4:**
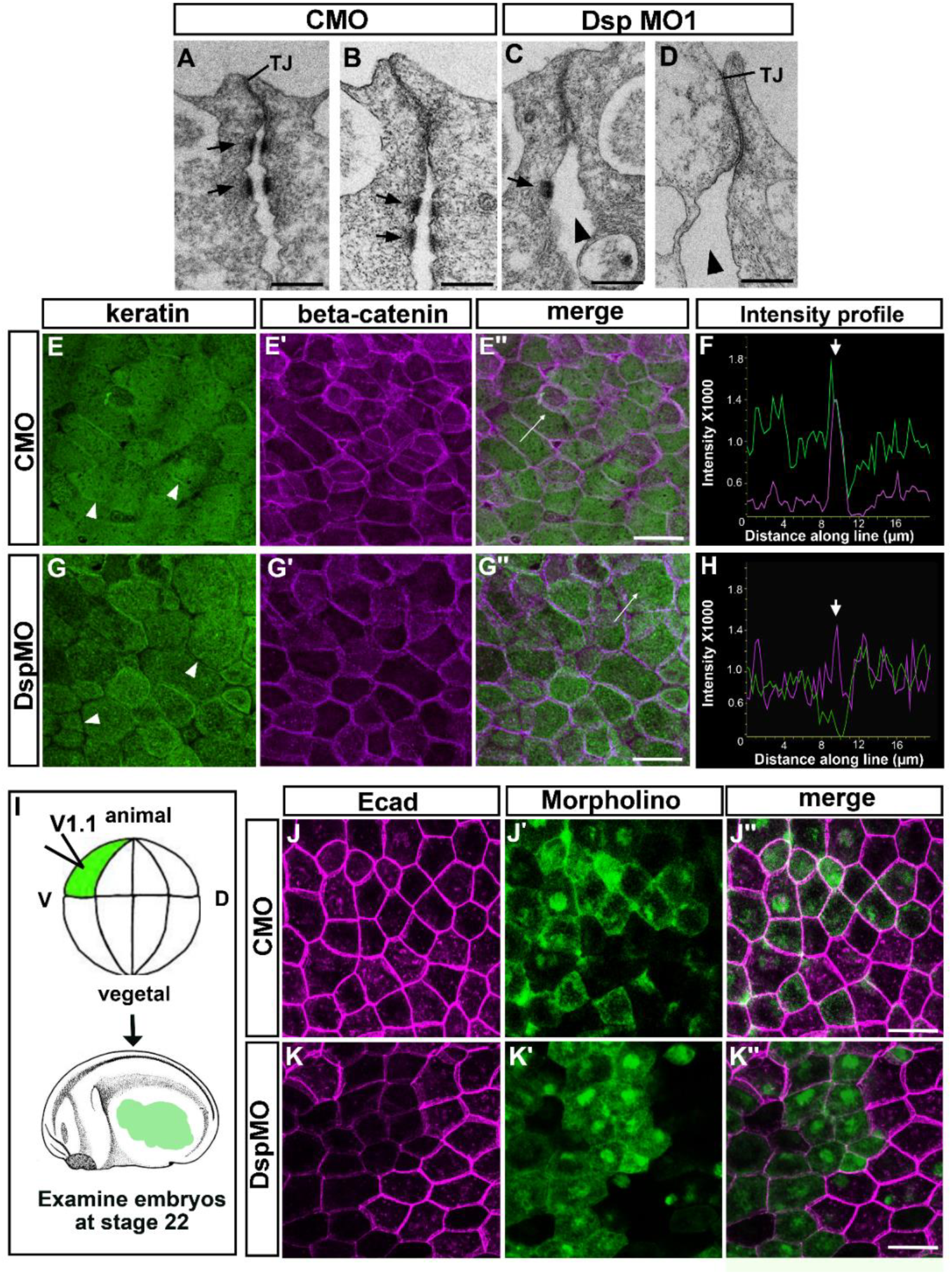
Decreased Desmoplakin results in changes in ultrastructure, keratin localization and E-cadherin levels. A-D) TEM images where black arrows indicate desmosomes located between outer epidermal cells. Arrowheads indicate a large space between cells with decreased desmosomes. Scale bars= 300nm. E-E‴) Representative control trunk epidermis labeled with keratin (E), beta-catenin (E′) and merge (E″). Scale bar = 25μm. F) Line graphs of the pixel intensities of keratin (green) and beta-catenin (pink) along the white line in E″. Arrow indicates highest beta-catenin intensity correlating with the cell-cell junction. G-G‴) Representative DspMO1 morphant trunk epidermis labeled with keratin (green). Beta-catenin labeling (pink) was used to provide a marker of the cortical region of the cell. (G), beta-catenin (G′) and merge (G″).Scale bar = 25μm. H) Line graphs of the pixel intensities of keratin (green) and beta-catenin (pink) along the white lines in G-G”. Arrow indicates highest beta-catenin intensity correlating with the cell-cell junction. I) Schematic showing targeted injection procedure. J-J″) Representative control trunk epidermis labeled with E-cadherin (J), showing the fluorescein labeled MO (J′) and merge (J″). Scale bar =25μm. K-K″) Representative control trunk epidermis labeled with E-cadherin (K), showing the Fluorescein labeled MO (K′) and merge (K″). Scale bar =25μm. Abbreviations= TJ=tight junction, Ecad=E-cadherin.

#### 3d) Dsp is required for normal keratin distribution

Desmosomes directly attach to the cytoplasmic network of keratins which is required for to maintain integrity and mechanical resilience across the epidermis. Thus, immunofluorescence was used to examine the distribution of keratins in Dsp morphants. Here whole embryo morphants were created by injection of morpholinos into the one-cell stage. In control embryos, epidermal keratin appeared as a relatively uniform network that extended to junctional beta-catenin labeling (Fig. 4E-E″). In some cells there also appeared to be an enrichment of keratin at the cell-cell interface (Fig 4E, arrowheads). Keratin intensity profiles were generated along an “intensity profile line” created between two cells (white lines with arrows in Fig. 4E″,G″). In the controls the profiles revealed that keratin intensity (green) was high adjacent to the high beta-catenin (pink and see arrow) at the cell-cell junction (Fig. 4F). In Dsp morphants there were gaps in keratin labeling adjacent to the junctional beta-catenin labeling in many cells (Fig. 4G-G″, arrowheads). Intensity profiles showed low keratin intensities adjacent to the high beta-catenin (arrow) consistent with cortical gaps (Fig. 4H). Such cortical gaps in keratin were also observed in the Crispr mutants and in DspMO2 morphants (Fig. S3).

These results suggest that in cells lacking adequate amounts of Dsp, the keratin is lower in the cortical region of the cell, where attachment to Dsp would take place. This lack of keratin attachment could affect mechanical resilience as well as a multitude of other functions.

#### 3e) Dsp deficiency results in decreased E-cadherin levels

Since epidermal integrity was so severely affected in Dsp morphants we next asked whether other junctions could defective. While, the EM images revealed intact tight junctions in Dsp morphants, it was more difficult to assess the adherens junctions. Therefore, a major adherens junction component, E-cadherin, was examined in Dsp morphants. However, we could not discern major differences in either beta-catenin or E-cadherin labeling when DspMO1 was injected into whole embryos due to the wide variability between embryos and samples (not shown). Therefore, we injected morpholinos into only one blastomere fated to become trunk epidermis (V1.1, 16 cell stage, Fig. 4I) so that morphant and normal cells could be directly compared side by side in the same embryo. Targeted control morpholinos did not result in any noticeable change in E-cadherin labeling in the trunk (Fig 4J-J″). However, a clear reduction in E-cadherin was observed in targeted Dsp morphants cells compared to their morpholino-free counterparts in the same embryo (Fig. 4K-K″). These results suggest that Dsp may be required for adherens junction strength and a decrease in this strength could contribute to the reduced mechanical integrity of the epidermis. In addition, changes in the adherens junction could also dramatically affect the actin cytoskeleton and cell-cell signaling during epidermal morphogenesis.

### 4. Epidermal morphogenesis is perturbed in Dsp morphants

We next asked whether the lack of Dsp in the epidermis specifically affected the morphogenesis and differentiation of this tissue. The *X. laevis* outer epidermal layer is comprised of several differentiated cell types including Multiciliated Cells (MCCs) and Small Secretory Cells (SSCs) that function to protect against bacterial infection. These cells are born in the inner epidermis and then move to their final location in the outer epidermis by radial intercalation. During this process the cell inserts itself at the vertex of 3-5 cells and then the apical surface expands. A combination of histology, immunofluorescence and cell tracking was utilized to assess the development of the epidermis including radially intercalating cells.

#### 4a) Decreased Dsp results in abnormal morphology of epidermal cells

General epidermal morphology was initially assessed in histological sections at stage 26-28 (hpf) when MCC cells are present in the outer epidermis. In the control embryos, the epidermis was comprised of an outer layer of cuboidal-like cells showing various phenotypes. For example, darker stained multiciliated cells (possibly MCCs) were interspersed between the larger lighter stained cells (Fig. 5A,B). The inner epidermis was comprised of flat, oblong cells that were stained dark blue (Fig. 5A, white arrow). A reduction of Dsp resulted in regions where the outer epidermal cells were less organized (Fig. 5B). For example, the large lighter stained cells were more globular in nature than the controls. The darker stained cells appeared “sunken” compared to the surrounding lighter stained cells (Fig. 5B, yellow arrows). Further multiple tufts of cilia was rarely observed on the surfaces of any cell. Also in Dsp morphants there appeared to be regions where there was more extracellular space underlying the inner epidermal cell layer (Fig. 5B, arrowheads). These observations demonstrate that a reduction in Dsp results in morphological perturbations in the embryonic epidermis.

**Figure 5:**
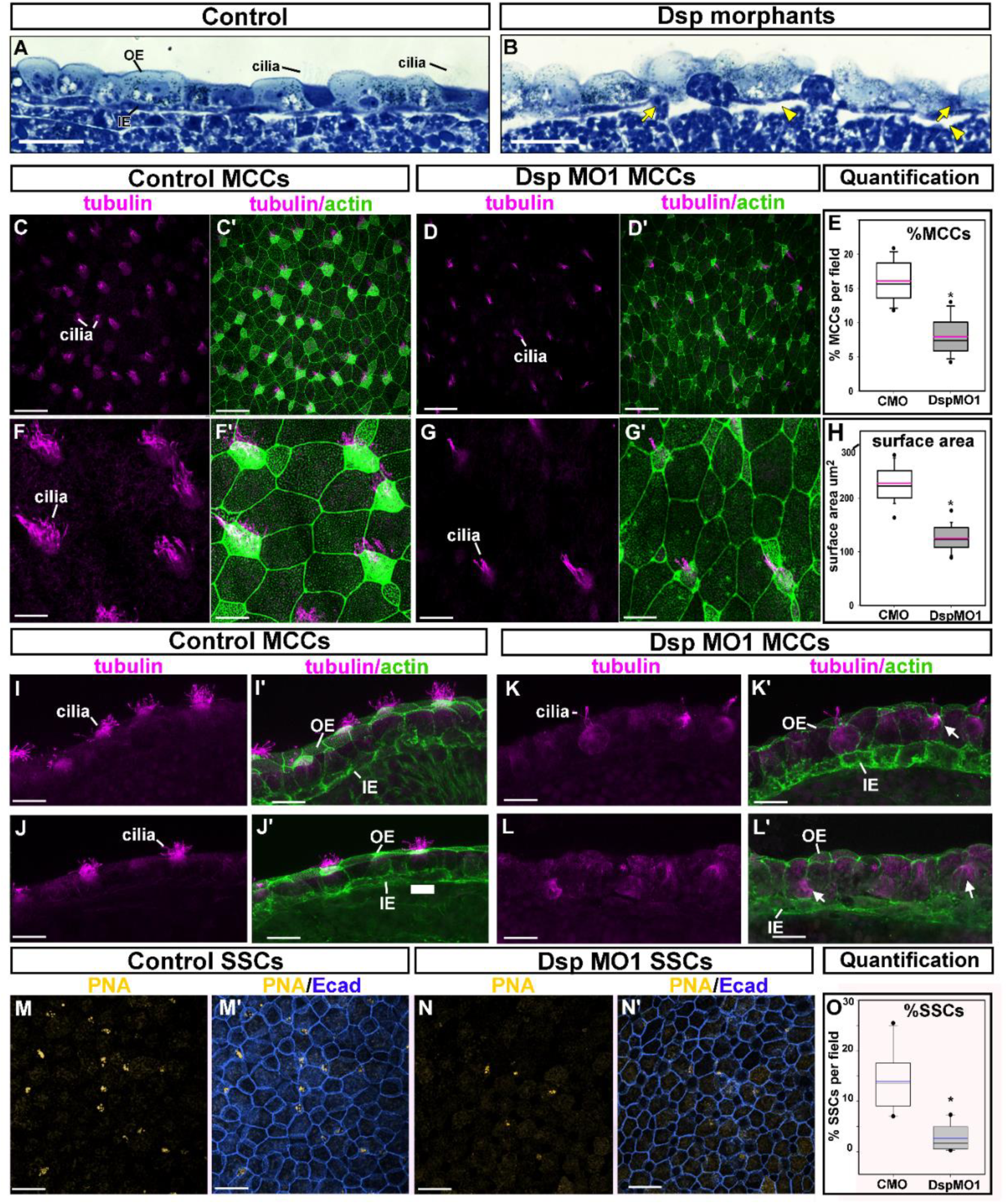
Decreased Desmoplakin results in abnormal epidermal morphology, MCC and SSC development. A-B) Histological sections of the epidermis in control (A) and DspMO1 morphants (B). Yellow arrowheads indicate space underlying the epidermis. Scale bars= 45μm. C-C′) Representative control trunk epidermis labeled with tubulin (C), phalloidin and tubulin (C′). Scale bars = 50μm. D-D′) Representative DspMO1 trunk epidermis labeled with tubulin (D), phalloidin and tubulin (D′). Scale bars = 50μm. E) Quantification of the percentage of tubulin positive cells in control compared to DspMO1 morphants. *=statistical significance. F-F′) Representative control trunk epidermis labeled with tubulin (F), phalloidin and tubulin (F′). 2X magnified region of C and C′. Scale bars =11μm. G-G′) Representative DspMO1 trunk epidermis labeled with tubulin (G), phalloidin and tubulin (G′). 2X magnified region of D and D′. Scale bar =11μm. H) Quantification of the surface area of tubulin positive cells in control compared to DspMO1 morphants. *=statistical significance. I-J′) Sections of two representative control trunk epidermis labeled with tubulin (I,J), phalloidin and tubulin (I′,J′). Scale bars = 22μm. K-L′) Sections of two representative DspMO1 trunk epidermis labeled with tubulin (J), phalloidin and tubulin (G′). Scale bar = 22μm. M-M′) Representative control trunk epidermis labeled with PNA (M), PNA and E-cadherin (M′). Scale bar =50μm. N-N′) Representative DspMO1 trunk epidermis labeled with PNA (N), PNA and E-cadherin (N). Scale bar =50μm. O) Quantification of the percentage of PNA positive cells in control compared to DspMO1 morphants. Blue line represents the mean. *=statistical significance. Abbreviations; PNA-peanut agglutinin, MCC=multiciliated cell, SSC=small secretory cell, OE=outer ectoderm, IE=inner ectoderm.

#### 4b) Decreased Dsp results in defects in MCC number, position and morphology

Our histological observations revealed epidermal defects that specifically included MCCs. These cells form multiple cilia on their apical surface that are identified here using alpha-tubulin labeling. Results demonstrated a 2.8 fold decrease in the relative number of alpha-tubulin positive patches on the surface of the epidermis of Dsp MO1 compared to controls (n=20, 2 experiments, p<0.0001, Fig. 5C-E).

In addition to a reduction in MCCs, it was also noted that Dsp MO1 morphants had a reduced cell surface often accompanied with fewer cilia (Fig. F-G′). Therefore, the surface area of the cells were measured using the phalloidin counterstain that marked the intense F-actin at the cell perimeter. Results indicated that indeed the DspMO1 morphants had an average 1.8 fold smaller MCC surface area when compared to the same cells in the controls (n=20, p<0.0001, Fig. 5F-H). In these experiments, we also noted that the apical phalloidin, was qualitatively less intense in the Dsp MO1 MCCs when compared to the controls (compare Fig. 5F′ and G′).

We next characterized the MCCs further in transverse sections of the epidermis labeled with alpha-tubulin and phalloidin. In control embryos, tubulin positive cilia was evident at the apical surfaces of cells of the outer ectoderm (Fig. 5I-J′). In the DspMO1 morphants where ciliated cells were observed, there were less cilia and the apical surface seemed to be sunken (Fig. 5 K,K′, arrows). Further, the apical enrichment of F-actin was not as apparent. In epidermal regions that seemed more severely affected, there was a less organized appearance of the inner and outer ectoderm (Fig. 5L,L′). Further, some tubulin enriched cells seemed to be located at the basal region of the outer ectoderm (Fig. 5L,L′ arrow). These observations are consistent with the histology reported above (section 4a).

To determine if the decreased numbers of MCCs was specific to the reduction in Dsp in the epidermis or whether it could be a more general non-specific effect, targeted injections to the trunk epidermis were performed as described above (section 3e and Fig. S4A). In control embryos, a uniform pattern of MCCs was apparent as indicated by alpha-tubulin labeling (Fig. S4B). However, MCCs were decreased in the DspMO1 positive regions (Fig. S4C,D) but not in adjacent regions with little or no morpholino present (Fig. S4C,D). These results suggest that indeed the loss of Dsp in the epidermis itself explain why there are defects in epidermal development. In addition, in these targeted injected embryos the epidermis was examined at stage 32 (40 hpf) and therefore any reduction we observed at earlier stages is not likely due to a delay in MCC development.

To determine whether other cells in the outer epidermis were also affected in DspMO morphants we next examined Small Secretory Cells (SSCs). These cells are also born in the inner epidermal layer and migrate to the outer epidermis by radial intercalation similar to the MCCs. SSCs were examined utilizing the marker peanut agglutinin (PNA) which is enriched in the secretory vesicles in these cells. Results reveled a 5.79 fold less cells with clear PNA positive accumulations in the epidermis of DspMO1 morphants compared to controls (n=20, p<0.0001, Fig. 5M-O). Together, these results suggest that Dsp is required for the development of at least two differentiated cell types in the epidermis, both of which undergo radial intercalation during their development.

#### 4c) Decreased Dsp results in defects in radial intercalation in the epidermis

Since the number and apical surface area of MCCs were affected in the Dsp morphants and mutants we hypothesized that Dsp is required for radial intercalation. To track radial intercalation, the epidermis was pulse labeled with biotin that adheres to the membranes of the outer ectodermal cells. Such labeling was performed from st.19 (21hpf) until st.20 (22hpf) during the first period of radial intercalation of primarily MCCs (Fig. 6A). Embryos that were fixed and labeled directly after this treatment demonstrated complete and relatively uniform biotin labeling of the epidermis (Fig. 6B). Biotin incubation was then followed by a 4 hour washout to allow unlabeled inner epidermal cells to emerge into the outer layer. Control embryos fixed after this washout had many biotin-negative regions (Fig. 6C, arrows) that were interspersed among biotin positive cells (Fig 6C, yellow labeling). The biotin-negative regions represented the cells that radially intercalated during the washout period. In the control embryos, the majority of such unlabeled biotin cells were also positive for alpha-tubulin, confirming that these cells indeed originated in the inner epidermal layer (Fig. 6C′-C″). Surprisingly, the DspMO1 morphants, had only a small reduction in the average number of unlabeled cells that was marginally significant (p=0.0475, n= 11, 3 experiments; Fig. 6D-H). The Box plot indicates a large amount of variability, where a subset of morphant embryos had the same or even more unlabeled cells than a subset of the controls (Fig. 6H). However, we also noted that in these embryos, especially those with high numbers of unlabeled cells, the size of the apical surfaces of such cells appeared smaller (Fig 6I-L). Thus, the surface area of the biotin-negative cells were quantified in the same images. Results revealed that indeed there was a statistically significant reduction in the average surface area of biotin-negative cells in the DspMO1 morphants when compared to controls (Mann Whitney, p=<0.0001, n=11, 3 experiments; Fig. 5M). A histogram analysis revealed that in controls the unlabeled surface areas followed a binomial distribution with a large proportion of cells greater than 100 μm^2^ (Fig 5N, see yellow line). On the other hand, in the Dsp MO1 morphants, a larger proportion of the unlabeled surface areas were less than 100 μm^2^ (Fig. 5O, see yellow line). These results reveal that DspMO1 morphants had more intercalating cells that had smaller apical surfaces. This result is consistent with the smaller apical surfaces of MCCs quantified in the previous section.

**Figure 6:**
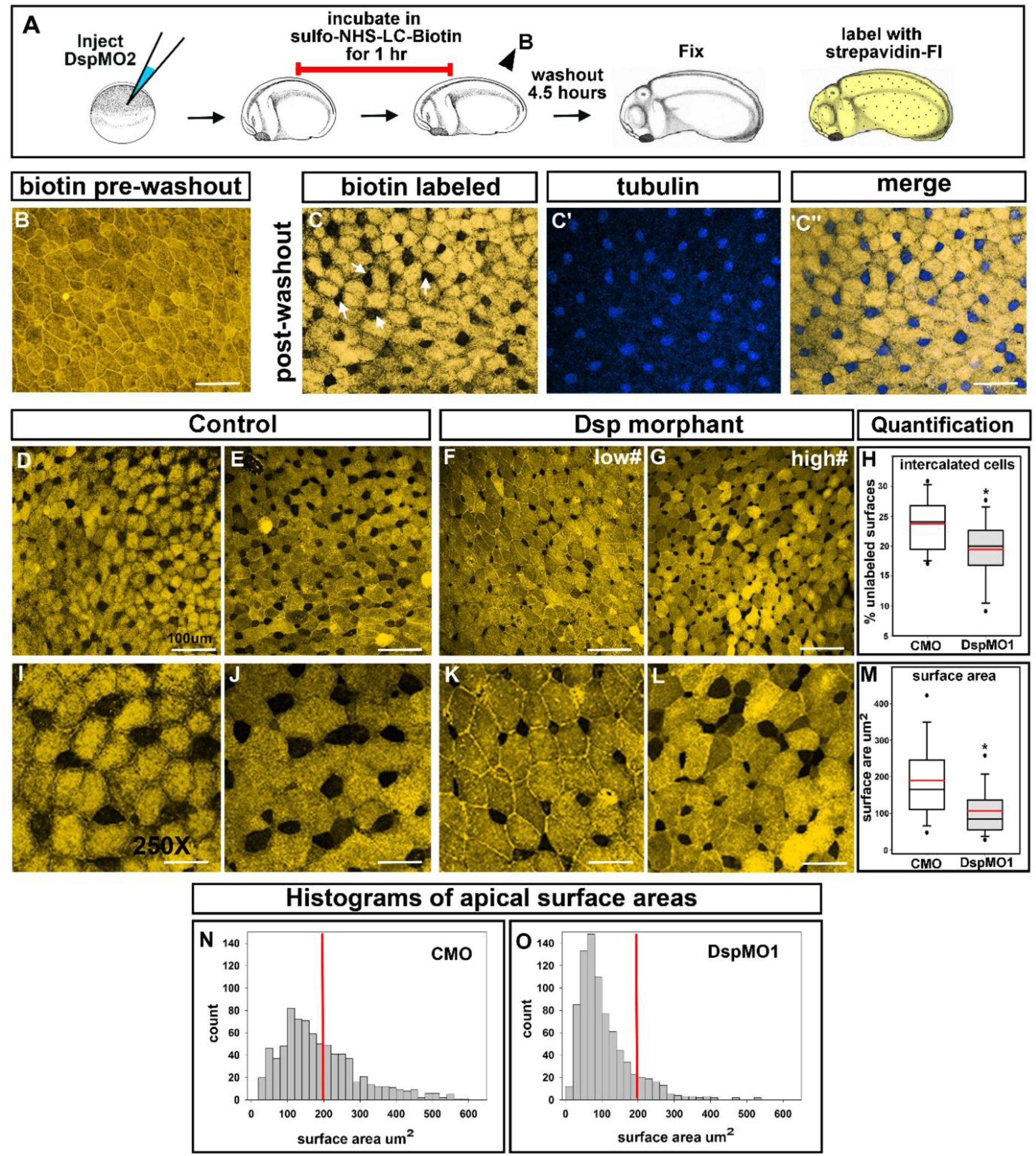
Tracking the epidermis. A) Schematic of outer epidermis labeling experiment. B) Representative image of trunk epidermis labeled with biotin directly after labeling without a washout period. Scale bar= 40μm. C-C″) Representative image of trunk epidermis double labeled with biotin (C) and tubulin (blue, C′) and merge (C″). Scale bar= 40μm. D-E) Two representative images of trunk epidermis from a control embryos after labeling with biotin. Scale bars=45μm. F-G) Two representative images of trunk epidermis from a DspMO1 morphant embryos after labeling with biotin. Scale bars =45μm. H) Quantification of the percentage of unlabeled cells in DspMO1 morphants compared to controls. Red line represents the mean. *= statistical significance. I-J) Two representative images of trunk epidermis from a control embryos after labeling with biotin at higher magnification. Scale bars=20μm. K-L) Two representative images of trunk epidermis from a DspMO1 morphant embryos after labeling with biotin at high magnification. Scale bars =20μm. M) Quantification of the relative change in surface area of unlabeled cells in DspMO1 morphants compared to controls. Red line represents the mean. *= statistical significance. N-O) Histogram analysis of surface areas of unlabeled cells in control (N) and DspMO1 (O). The red line indicates the surface area of 200μm^2^ for comparisons.

Taken together, our results reveal the requirement for Dsp in radial intercalation, most notably in the process of apical expansion.

### 5. Expression of a mutant Dsp in *X. laevis* mimics knockdown of Dsp

We next tested the specificity of Dsp in embryonic development and MCC formation by expressing a mutant form of the human Dsp (DP-NTP). This construct is missing the C terminal domains which impair interactions with intermediate filaments (Fig. S5A). However, the N-terminus is present in DP-NTP and therefore desmosomal cadherin associations and cell-cell adhesions are maintained. In *X. laevis* outer epidermis, we confirmed that DP-NTP displayed membrane association and replaced endogenous Dsp (Fig. S5B-B″, see arrow for example). In addition, keratin was reduced in the cortical regions of cells expressing DP-NTP suggesting that indeed Dsp-keratin interactions were impaired (Fig. S5C,D). Further, this phenotype mimics our observations of keratin localization in Dsp morphants and mutants. Embryos expressing DP-NTP were smaller in size and displayed hyperpigmentation similar to Dsp morphants and mutants (Fig. S5E,F). Finally, DP-NTP expressing embryos also displayed a reduction in MCCs (Fig. S5G,I′). These results indicate that many of the perturbations noted in embryos deficient in Dsp are likely specific and could be due to the loss of connections to the cytoskeleton.

### 6. Dsp is required for the formation of structures that also use intercalation type cell movements during their development

Ectodermally derived structures that use intercalation type cell movements during their formation include the embryonic mouth, eye and fin. Therefore, we next sought to determine whether Dsp is required in the morphogenesis of these structures. We targeted Dsp morpholinos to these structures using specific blastomere injections to rule out secondary effects from gastrulation or heart defects. DspMO1 was injected into a dorsal blastomere (D1.2; Xenbase) at the 16 cell stage that fate maps to the face as well as the eye (Fig. 7A). In these embryos, the injected side of the embryo showed a failure of the embryonic mouth to form in 100% of the embryos (n=20, 2 experiments; Fig. 7B-C″). In addition, eye defects including a small eye and a protruding lens were also observed in 90% of the DspMO1 morphants (n=17, 2 experiments; Fig 7D-F″). In embryos where a ventral blastomere (V.2.2, Xenbase) at the 16 cell stage was targeted, the fin was labeled (Fig. 7G). In these partial morphants the fin was ruffled, lacking the smooth appearance of the controls (n=23, 2 experiments; Fig. 7H-I″). These results demonstrate the requirement of Dsp for mouth, eye and fin development. Importantly, we suggest the possibility that Dsp could be required more generally during morphogenesis of structures utilizing intercalation type cell movements.

**Figure 7:**
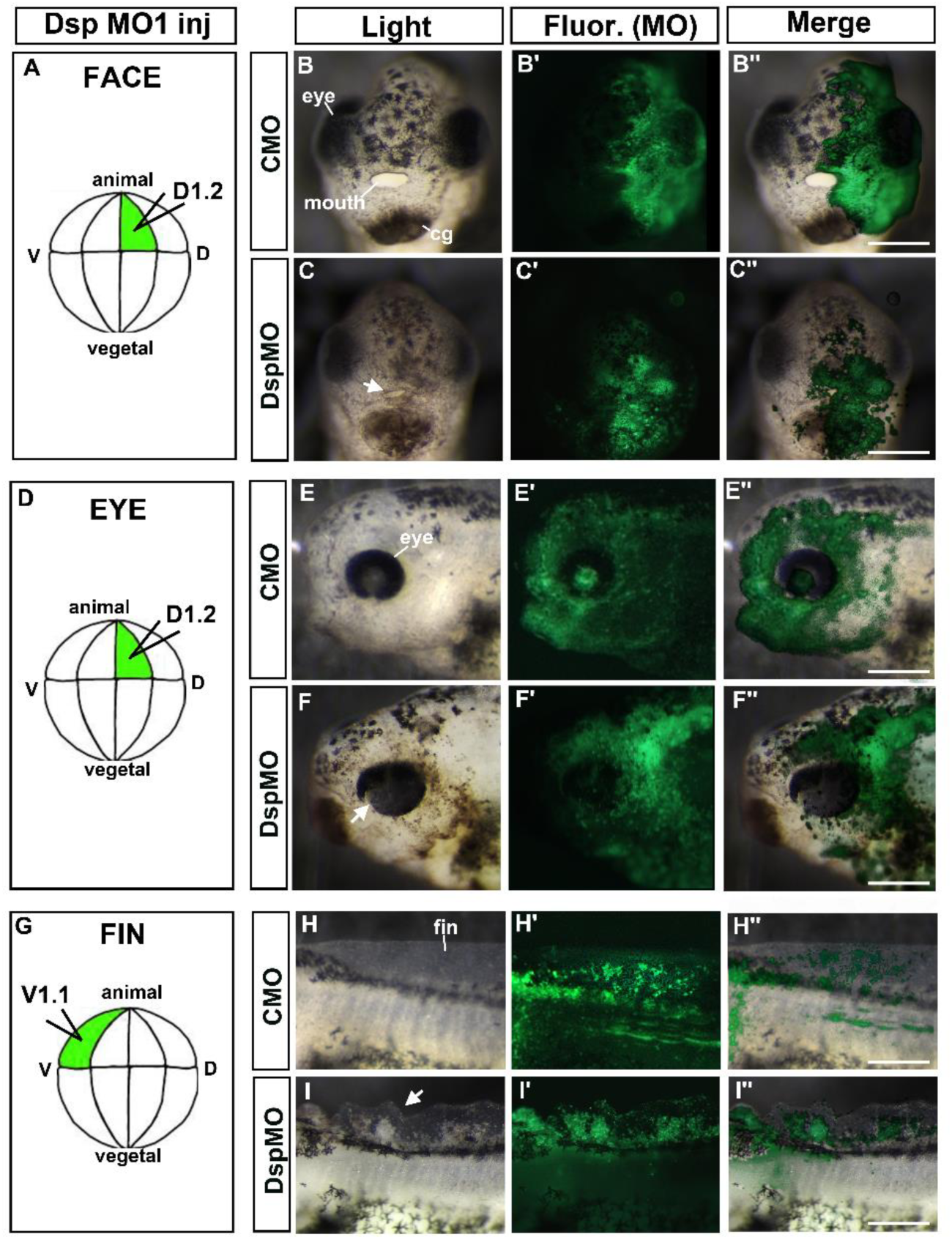
DspMO1 targeted to the face, eye and fin causes developmental defects. All scale bars= 85μm. A) Schematic showing the blastomere injected with MO to target the face. B-B″) Frontal view of the face of a representative embryo with control MO in the face. Shows an embryo with light microscope (B), fluorescent (B′) and merge (B″). The embryonic mouth forms normally. C-C″) Frontal view of the face of a representative embryo with DspMO1 in the face. Shows embryo with light microscope (C), fluorescent (C′) and merge (C″). The embryonic mouth does not form (arrow). D) Schematic showing the blastomere injected with MO to target the eye. E-E″) Lateral view of the head of a representative embryo with control MO in the eye. Shows an embryo with light microscope (E), fluorescent (E′) and merge (E″). F-F″) Lateral view of the head of a representative embryo with DspMO1 in the eye, shows embryo with light microscope (F), fluorescent (F′) and merge (F″). The abnormal eye is indicated by an arrow. G) Schematic showing the blastomere injected with MO to target the dorsal fin. H-H″) Lateral view of the head of a representative embryo with control MO in the dorsal fin. Shows an embryo with light microscope (H), fluorescent (H′) and merge (H″). I-I″) Lateral view of the head of a representative embryo with DspMO1 in the fin, shows embryo with light microscope (I), fluorescent (I′) and merge (I″). The abnormal fin is indicated by an arrow.

## Discussion

### The structure and adhesive functions of desmoplakin are conserved in the embryonic epidermis of X. laevis

Desmosomes are especially distinct by TEM imaging, where they appear as electron-dense structures consisting of two identical cytoplasmic plaques and a central core that spans the intercellular space (reviewed in (Berika and Garrod, 2014; Garrod and Chidgey, 2008)). This ultrastructure is observed in all vertebrates examined to date and now we have included *X. laevis* embryos (Borysenko and Revel, 1973; Farquhar and Palade, 1965; Fleming et al., 1991; Garrod and Chidgey, 2008; Goonesinghe et al., 2012; Holbrook and Odland, 1975). Desmoplakin, a unique and critical protein member of the desmosome, is also highly similar in both sequence and composition of the functional domains in vertebrates. *X. laevis* desmoplakin is no exception. These structural similarities suggest the strong possibility that the desmosome and, desmoplakin in particular, have well conserved roles across vertebrates including *X. laevis* embryos. Certainly our experiments demonstrate that desmoplakin does indeed have some conserved functions in *X. laevis* embryos. For example, one of the most well-known roles for desmosomes in mammals is to protect the epidermis from mechanical stress. Humans and mice with mutations in Dsp and other desmosomal genes have skin fragility (Smith et al., 2012; Whittock et al., 1999; Whittock et al., 2002). Similarly in *X. laevis* embryos, a reduction in Dsp resulted in skin fragility and reduced resistance to mechanical stresses. The epidermis became damaged upon impact and sheer forces more readily than controls. Thus skin fragility, possibly due to decreased cell adhesion between cells of the epidermis, is a common defect in vertebrates (including *X. laevis* embryos) lacking Dsp. Another common feature of humans with desmosomal diseases is skin blistering. One possible reason for this problem is that the epidermal layers do not adhere to each other and this results in fluid filled spaces between such layers. In agreement with this idea, separations between skin layers, specifically the basal and spinous layers were reported in mice with dsp mutations (Gallicano et al., 2001; Vasioukhin et al., 2001). In *X. laevis*, we also observed similar spaces between embryonic tissue layers and a percentage of Dsp morphants and mutants also had blister like formations. Together, these data demonstrate a role for desmosomes in maintaining overall epidermal connectivity not just within a skin layer but also between different layers in vertebrates, *X. laevis* embryos included. Since these fundamental roles of desmosomes are conserved in *X. laevis* then we might predict that this model could be a useful tool to uncover additional roles during embryogenesis.

### Dsp is required for cytoskeletal and adherens junction components

In the epidermis, desmoplakin connects to the cytoskeletal network through keratins. For example, in humans, mutations in the dsp gene is associated with reduced keratin attachment to the desmosome (Favre et al., 2018). In mice, where dsp has been specifically deleted in the epidermis, the keratin network does not extend to the periphery or cortical region of the cells (Vasioukhin et al., 2001). A similar change in the distribution of keratin was also observed in desmosomal deficient cells in vitro (Vasioukhin et al., 2001). We too observed a reduction of keratin at the epidermal cell-cell boundaries in *X. laevis* Dsp morphant embryos. This alteration of keratin distribution could be expected to have dramatic effects during development especially since keratins are important for cell morphology and migration (Nishimura et al., 2018; Seltmann et al., 2013). For example, loss of keratin in the embryo results in misdirected lamellapodial protrusions and disorganized actin networks that translate into defects in tissue level collective migration (Sonavane et al., 2017). Thus, it is likely that keratins integrate the cell mechanics necessary for morphogenetic movements (Klymkowsky et al., 1992).

In addition to keratins, loss of Dsp affected other major components of the cell’s architecture in *X. laevis*, namely the adherens junctions and the associated actin cytoskeleton. Adherens junctions, like desmosomes, use cadherins (such as E- or N-cadherin) to bridge the extracellular space and connect cells to each other. The intercellular domain of cadherins bind to catenins which, together with other accessory proteins, link the junction to actin filaments (reviewed in (Yonemura, 2011)). In Dsp morphants there is reduced e-cadherin associated with the adherens junctions in the developing epidermis. Further, we also noted a decrease in the apical accumulation of actin in multiciliated epidermal cells. This is consistent with reports in mice where conditional knockout of Dsp in the epidermis also resulted in a decrease in adherens junctions in some skin layers as well as disorganized actin cytoskeleton (Vasioukhin et al., 2001). In addition, mouse epithelium deficient in desmoplakin, had decreased actin stability and altered microvillar structure (Sumigray and Lechler, 2012). Together these results indicate possible crosstalk between desmosomes and its sister junction, the adherens junction, in the developing embryo. Such crosstalk has certainly been shown in cultured cells. For example, adherens junction associated proteins, e-cadherin and actin are both required for desmosome assembly (Lowndes et al., 2014; Shafraz et al., 2018). Further, disruption of the desmosomal protein, Desmoglein 3, decreased e-cadherin at the junction and perturbed actin organization in cultured epithelial cells (Moftah et al., 2017). Therefore, our results and the work of others demonstrate that disruption of the desmosome can affect the adherens junctions and associated cytoskeleton. Since, adherens junction-actin complexes direct processes such as cell shape changes and migration (Harris and Tepass, 2010; Nishimura and Takeichi, 2009) then not surprising is our results where desmosome disruption affected morphogenesis in the developing embryo.

### Dsp is required for morphogenesis of the epidermis in X. laevis

A major cellular process in epidermal morphogenesis is radial intercalation where the inner cell layer migrates to a position in the outer cell layer. Here we have uncovered a requirement for Dsp in the radial intercalation of epidermal cells. Pulse labeling the outer epidermis of Dsp morphants just as MCCs are intercalating, resulted in emerging cells with smaller surface areas. Further, newly emerged cells appear to be enriched in apical Dsp during early epidermal development. Correlated to the decrease in apical surface size and a reduction in intercalating cells, was a decrease in the number of MCCs. The MCCs that did differentiate had smaller surface areas, appeared less ciliated and had decreased apical actin. These results suggest a role for Dsp in the last step of intercalation of MCCs, apical expansion (see model Fig. 8).

**Figure 8:**
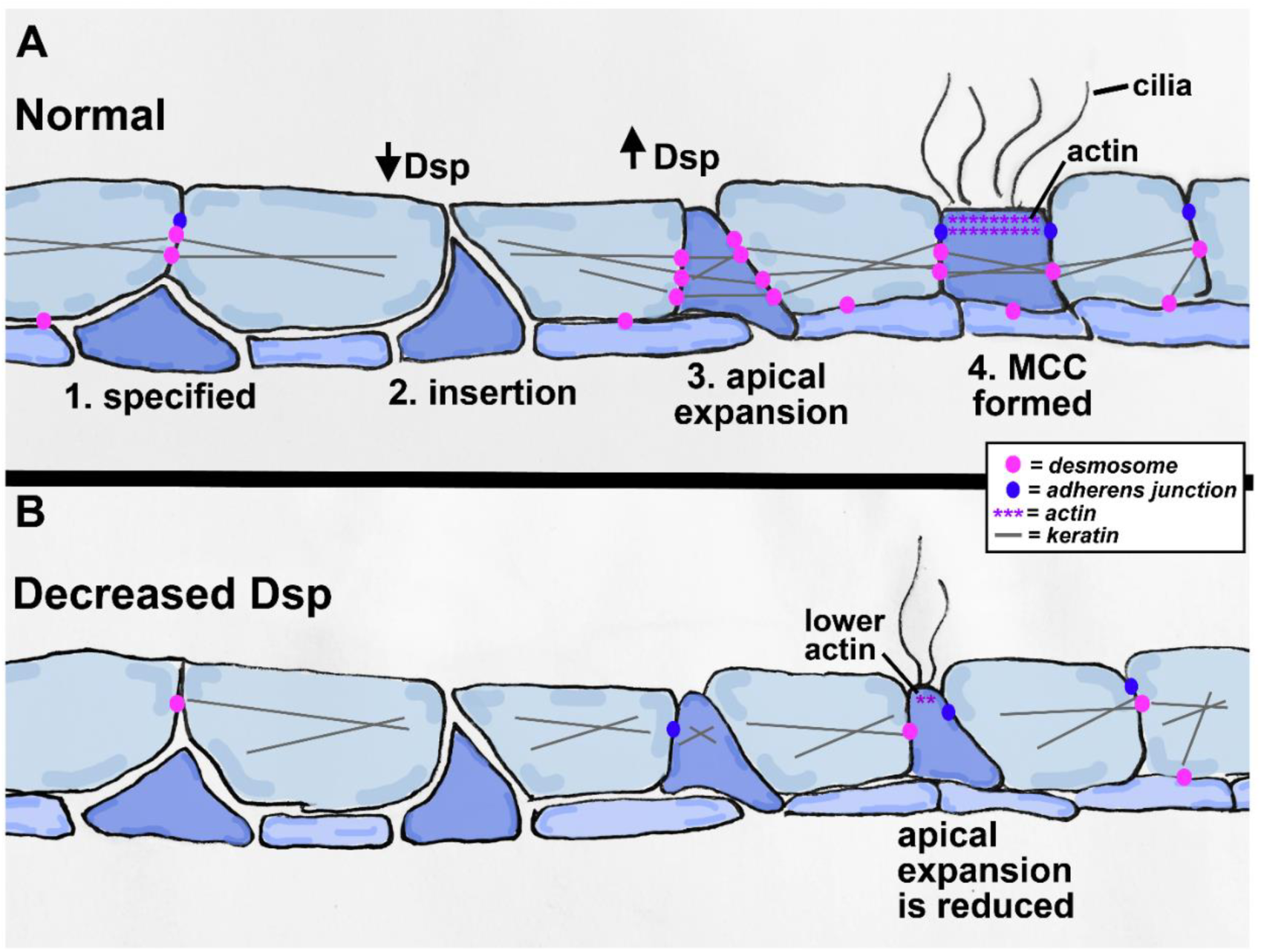
Model of radial intercalation of the epidermis in normal embryos (A) and those with reduced Dsp (B). Dsp (pink) connects with keratins and is increased during apical expansion. With reduced Dsp, the intercalating cell does not have properly localized keratins or apical accumulation of actin and thus the apical surface does not expand sufficiently.

Dsp together with keratins may have a direct role in apical expansion by providing the emerging cells and with the rigidity needed to support apical expansion. Certainly, desmosomes themselves have been shown to provide mechanical strength to cells (Baddam et al., 2018; Puzzi et al., 2018) and keratins convey stiffness to cells important for cell shape changes (Ramms et al., 2013). Dsp and keratin attachment could also be required to provide adequate tension in the field of outer epidermal cells. Such tension is predicted to be necessary to support cell shape changes in the emerging cells (Sedzinski et al., 2016). Dsp could also have an indirect role in apical expansion, where disruption of Dsp affects actin accumulation. Rho directed apical accumulation of actin has been shown to be critical for apical emergence of intercalating cells (Sedzinski et al., 2016, 2017). In support of this idea is evidence that desmosome associated plakophilin 2 recruits active RhoA to drive actin reorganization (Godsel et al., 2010). Disruption of Dsp could also affect other cell signaling pathways and transcriptional modulation of genes which could be necessary for apical expansion. For example, decreased Dsp could alter plakoglobin localization and in turn alter Wnt/beta-catenin signaling (Martin et al., 2009; Yang et al., 2012), a pathway required for a multitude of downstream targets (for example see (Kjolby and Harland, 2017)). We have also demonstrated that the role of desmoplakin is not likely specific to MCC’s or to a specific time in development. In addition to a reduction in MCCs, Dsp morphants had less small secretory cells, an epidermal cell type that also intercalate from the inner cell layer later in development. These results suggest that Dsp and the desmosome are required in a general rather than cell type specific process. Future experiments will be essential to further dissect the role of Dsp and the desmosome in epidermal morphogenesis.

### Dsp is required for the morphogenesis of ectodermally derived structures

In addition to epidermal radial intercalation, there are other processes that require similar types of cell movements in ectodermally derived tissues and that also express desmoplakin. For example, early in development, intercalation type cell movements are required for convergent extension during gastrulation (reviewed in (Walck-Shannon and Hardin, 2014)). Consistent with this is that zebrafish injected with translation blocking morpholinos targeting desmosomal cadherins have convergent extension defects (Goonesinghe et al., 2012). Dsp knockout mice, did proceed through gastrulation but were extremely small with head and neural defects characteristic of convergent extension problems (Gallicano et al., 2001). Our Dsp morphant or mosaic dsp mutant *X. laevis* embryos were shorter, indicative of convergent extension deficiencies. However, otherwise embryos proceeded through gastrulation relatively normally without major head defects. This could be explained by our use of splice blocking Dsp morpholinos that only target zygotic expression of Dsp. We know that Dsp mRNA is maternally loaded and therefore suspect that these maternal transcripts provide enough Dsp to allow embryos to develop through gastrulation relatively normally. Additionally, mosaic Dsp mutants could also have enough normal cells, with mutant Dsp cells interspersed, to prevent dramatic gastrulation defects. Further, combining splice blocking morpholinos with targeted blastomere injections has allowed us to investigate roles of Dsp without large scale interruption of gastrulation. In doing so we were able to observe defects in ectodermally derived structures such as the mouth, eye and fin that develop much later in development. These structures also utilize intercalation type cell movements during their development. For example, intercalation of cells in the developing embryonic mouth is necessary for eventual perforation of the structure (Dickinson and Sive, 2006). Embryos with deficiency of Dsp in the developing face did not form a perforated embryonic mouth. Interestingly, humans with birth defects caused by mutations to desmosomal proteins can present with a number of craniofacial abnormalities involving the tongue and teeth which could be rooted in problems with early orofacial development (reviewed in (Celentano and Cirillo, 2017; Celentano et al., 2017). Also, similar to the epidermis, primary lens cells move from one layer to another to form secondary lens cells that contribute to lens growth (reviewed in (Cvekl and Ashery-Padan, 2014)). Defects in lens development can also affect retina morphogenesis (Fuhrmann, 2010). *X. laevis* embryos deficient in Dsp had both lens and retinal defects. Humans with epidermolysis bullosa, which can be caused by a mutation in the dsp gene, have a high likely hood to have ophthalmic complications (McDonnell et al., 1989; Tong et al., 1999). Finally, during dorsal fin development, epidermal cells on each side of the body fuse and integrate to form one layer of cells (Tucker and Slack, 2004). This process closely resembles radial intercalation. *X. laevis* embryos with Dsp targeted to the dorsal ectoderm had a ruffled abnormal dorsal fin. These results suggest the possibility that desmosomes are important for intercalation type movements in forming epidermal structures. Certainly it would be interesting to investigate this hypothesis further in mammals where hair placodes also use cell intercalation to form and hair abnormalities are one of the most common defects associated with desmosomal disorders.

Together, our results suggest that desmosomes and desmoplakin in particular could be important for more that mechanical resilience, but also integral for directing cytoskeletal changes necessary for a broad range of cell movements in the developing embryo. Further, *X. laevis* could prove to be a valuable tool to begin to uncover the mechanisms by which desmosomes are involved in morphogenesis which in turn could help explain why particular defects occur in humans with desmosomal defects as well as in the development of therapeutic options.

## Acknowledgements

We would like to thank Daniel Conway (VCU, Department of Biomedical Engineering) for providing us with a modified DP-NTP construct and for his insightful contributions to the study. In addition, we would like to thank Jakub Sedzinski, Greg Walsh, James Lister, and Rita Shiang whose ideas and thoughts also helped shape this work. We thank the VCU Microscopy Facility, supported, in part, by funding from NIH-NCI Cancer Center Support Grant P30 CA016059 for helping with our TEM preparation, training and imaging as well as the VCU Biology Microscopy Core for providing confocal use. We thank VCU undergraduate students Morgan Van Driest and Skyler Kuhn for their help with comparative bioinformatics analysis of Dsp. We especially thank Deborah Howton for helping with the frog care essential for this study. We apologize to those whose work could not be cited here due to space constraints.

